# Social transmission of valence-linked new knowledge without firsthand experience in mice

**DOI:** 10.1101/2023.08.27.555038

**Authors:** Ryang Kim, Haruhiko Bito, Takashi Kitamura

**Affiliations:** Department of Neurochemistry, Graduate School of Medicine, The University of Tokyo, Tokyo, 113-0033, Japan; Department of Psychiatry, University of Texas Southwestern Medical Center, Dallas, TX 75390, USA; Department of Neuroscience, University of Texas Southwestern Medical Center, Dallas, TX 75390, USA

## Abstract

Animals can acquire new knowledge by observing others’ contexts and behavior, a process called social learning, which is essential for animals to survive in nature. While the social transmission of food preference (STFP) was previously adopted as a social learning test, several studies pointed out that non-social parameters might influence its food preference. We re-evaluated critical steps in the STFP test and designed an improved social learning test, which we now refer to as social transmission of food finding (STFF). A significant advance in the STFF test is the demonstration that mice learned the odor-food relationship with either positive or negative valence from the demonstrator without prior experience in the observer. Furthermore, a circuit dissection experiment showed that hippocampal function was differentially involved during learning and recall for STFF. Thus, STFF constitutes a highly advantageous social learning model in which valence-linked new knowledge can be socially transmitted without firsthand experience.

## Introduction

Animals can associate aversive or appetitive stimuli with cues that predict future outcomes, which is crucial for survival in nature. Given that direct experience entails significant risk and effort, there is a considerable evolutionary incentive for social learning among various animal species^1–4^. In rodents, both appetitive and aversive types of social learning tests have been developed. For example, observational fear conditioning in rodents^5–8^ is a vicarious associative fear learning, allowing the observer to ascertain dangerous stimuli and situations by witnessing the demonstrator’s aversive moments. The social transmission of food preference (STFP) has been demonstrated for a long time as a reward-based spatial-independent social learning test^3,9,10^. This test enables observers to learn that an unknown odor is associated with food by smelling it from the demonstrator that has consumed the odor-associated food without directly observing the demonstrator’s feeding behavior^11^. Once learned from the demonstrator, the observer exhibits a “preference” for the food with the smelled odor compared to the same food with a novel odor. However, the STFP test often includes the habituation session with a food-filled cup, which allows the observer to directly learn about the presence of food in the cup, potentially influencing food preference. Furthermore, previous studies have also raised concerns about the STFP that an odor may be sufficient to generate a choice to a food with the same odor^12,13^. These issues raised the question that the STFP test may have a limitation for interpreting data as social learning. In this study, we first confirmed that the food preference observed in the STFP test could be influenced by non-social components, as previously reported^12,13^. Therefore, we next designed an improved social learning test in mice, named social transmission of food finding (STFF) and we examined whether animals could acquire new knowledge from the demonstrator without firsthand experience. In the STFF test, we excluded the habituation session from the protocol, and we evaluated whether the observer acquired knowledge about an unknown odor associated with food from the demonstrator by measuring the animal’s latency to eat the food in the cup. After multiple interaction sessions, we found that the latency to eat the food with this associated odor was shorter compared to the control group during the test session. The habituation with the odor itself without the presence of the demonstrator did not facilitate the latency to eat a portion of food, indicating that the STFF is a social learning test. Importantly, in the STFF test, the observer recognized positive and negative valence with an unknown odor in a manner dependent on the demonstrator’s state (healthy or sick). These results conclude that the STFF is a highly advantageous valence-rich social learning model. Finally, we examined the roles of dorsal hippocampal function in the STFF and found that hippocampal function was differently involved during the learning and recall processes for STFF.

## Results

### Learning about a new food significantly facilitates eating behavior upon re-exposure

Typically, animals do not immediately try to eat an unknown object presented in front of them, even if it is edible, through a phenomenon called food neophobia^14,15^. We speculated that if animals somehow learned that a previously unknown object is related to food, they might start eating it promptly under food-deprived conditions. To test this, mice were subjected to two types of tests after a food deprivation period (see methods) (Fig. 1a-f). In the first experiment, mice were exposed to a novel feeding jar filled with a novel powder food or a familiar food (Fig. 1a). The group of mice exposed to the familiar food-filled feeding jar (Familiar) showed a significantly reduced latency to start eating compared to the group of mice exposed to the novel powder food-filled feeding jar (Novel) (Fig. 1b). Additionally, the familiar group showed significantly more food consumption than the novel group (Fig. 1c). In the second experiment, one group of mice was exposed to the novel powder food-filled jar during the habituation session (with group, Fig. 1d), while the other group of mice was not (without group, Fig. 1d). Then both groups of mice were exposed the same powder food-filled jar during the test session. The latency to start eating the food in the jar and the volume of food consumption were significantly faster and larger for the with group as compared with the without group, respectively (Fig. 1e-f). These data suggest that prior knowledge about the food induces a reduced latency to start eating accompanied by larger food consumption, even when the food is in a novel feeding jar for the mice. These results indicate that the latency to start eating and the volume of food consumption can be reliable behavioral indicators to test if mice could learn that the novel odor is food-related.

**Fig. 1:**
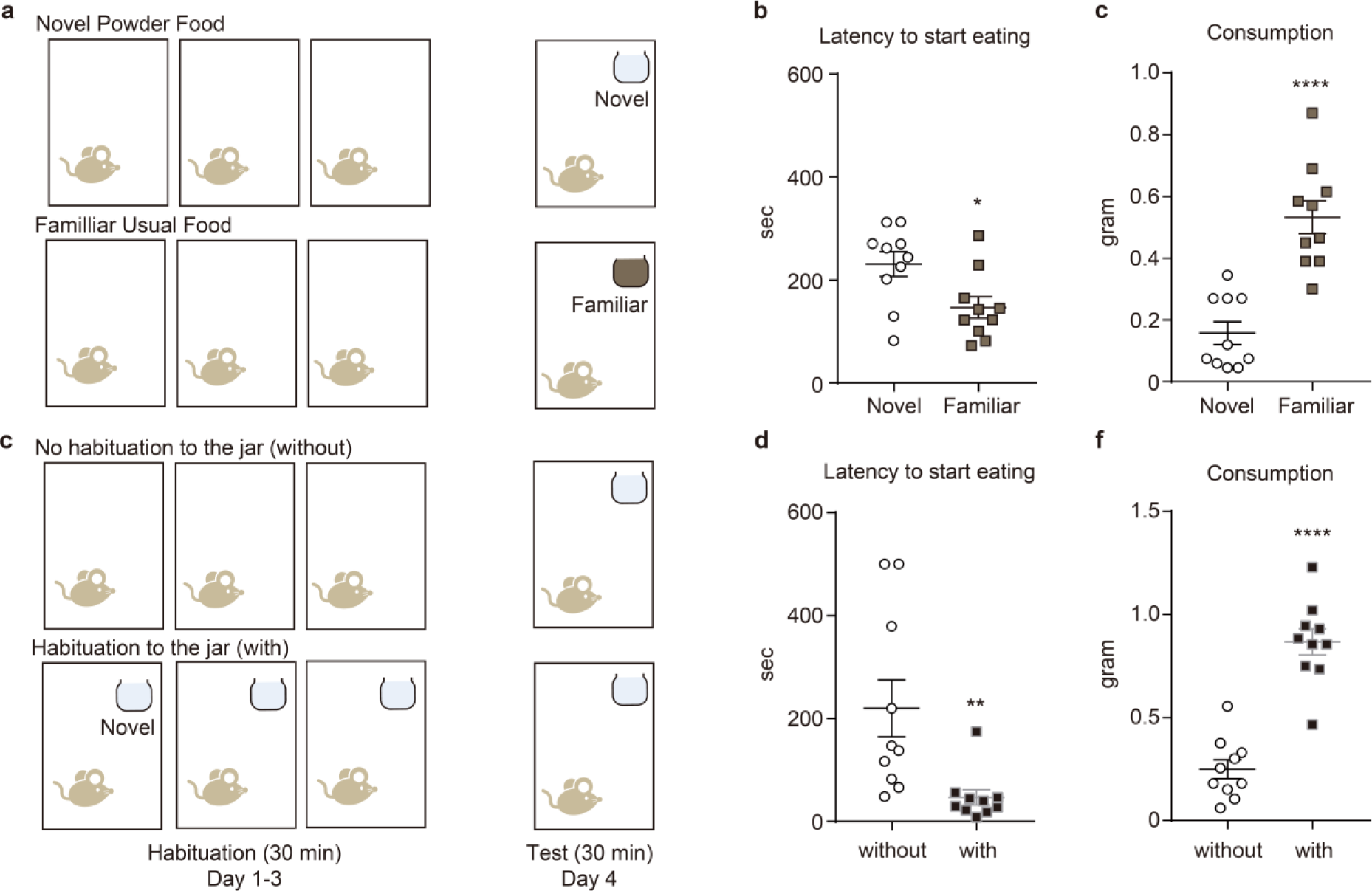
Learning about a new food significantly facilitates eating behavior upon re- exposure. **(a)** Schematic illustration of the protocol. **(b)** Latency to star eating the food in the jar during the test session (n = 10. Two-tailed unpaired *t*-test, t_18_ = 2.661, *P = 0.0159). **(c)** Consumption of the food during the test session (n = 10. Two-tailed *t*-test, t_18_ = 5.781, ***P < 0.0001). **(d-f)** Analysis of feeding behavior when presented with or without a novel-food filled- jar during the habituation session. **(d)** Schematic illustration of the protocol. **(e)** Latency to start eating the food in the jar during the test session (n = 10. Two-tailed unpaired *t*-test, t_18_ = 3.012, **P = 0.0075). **(f)** Consumption of the food during the test session (n = 10. Two-tailed unpaired *t*-test, t_18_ = 7.884, ****P < 0.0001).

### Food preference in the STFP test is driven by non-social effects

Next, we attempted to determine the critical factor driving food preference in an STFP test. The STFP test in mice consists of three sessions: habituation, interaction, and test sessions (Fig. 2b)^9,16,17^. During the habituation session, the demonstrator group and the observer group of mice were placed in a new cage with a novel powder food-filled in novel jars for 30 min for three consecutive days. During the interaction session, the observer mouse interacted with the demonstrator mouse that had consumed cumin-odorized food before interaction with the observer, thereby allowing the observer to smell the cumin odor from the demonstrator. We used cumin as the cued odor for statistical reasons (Fig. 2b)^9^ because mice showed a natural preference for vanillin-odorized food over cumin-odorized food (Fig. 2a). One day after the interaction session, the observer animal was subjected to the test session in which both cumin- and vanillin-odorized food-filled jars were placed in the cage. The cued mice group consumed significantly more cumin-odorized food than the non-cued control group, which interacted with the demonstrator that had consumed plain, non-odorized food (Fig. 2c). We also found the total consumption of these foods unchanged between two groups (Fig. 2c). These results were previously interpreted as showing that mice *socially* learned a safe food odor through the smell of the demonstrator. To test if the preference is social-dependent, we used a cumin-odorized cotton ball, instead of the demonstrator, during the interaction session. Here, we found that the mice that directly or indirectly interacted with the cotton ball showed a similar preference for the odor even when it was exposed in the absence of social interaction (Fig. 2d-g). This indicated that an odor itself contains sufficient information to generate a preference without necessitating any interaction with the demonstrator, in keeping with some previous reports^12,13^. Furthermore, any preference was lost when the observer mice did not experience the habituation session to learn that the jar is associated with food and were then directly exposed to the cumin-odorized cotton ball in the interaction session (Fig. 2h-i)^18,19^. This raised the possibility that the observed food preference in the data above was simply triggered by the co-incidence of jar-food association memory and the familiarity with the cumin odor during the test session.

**Fig. 2:**
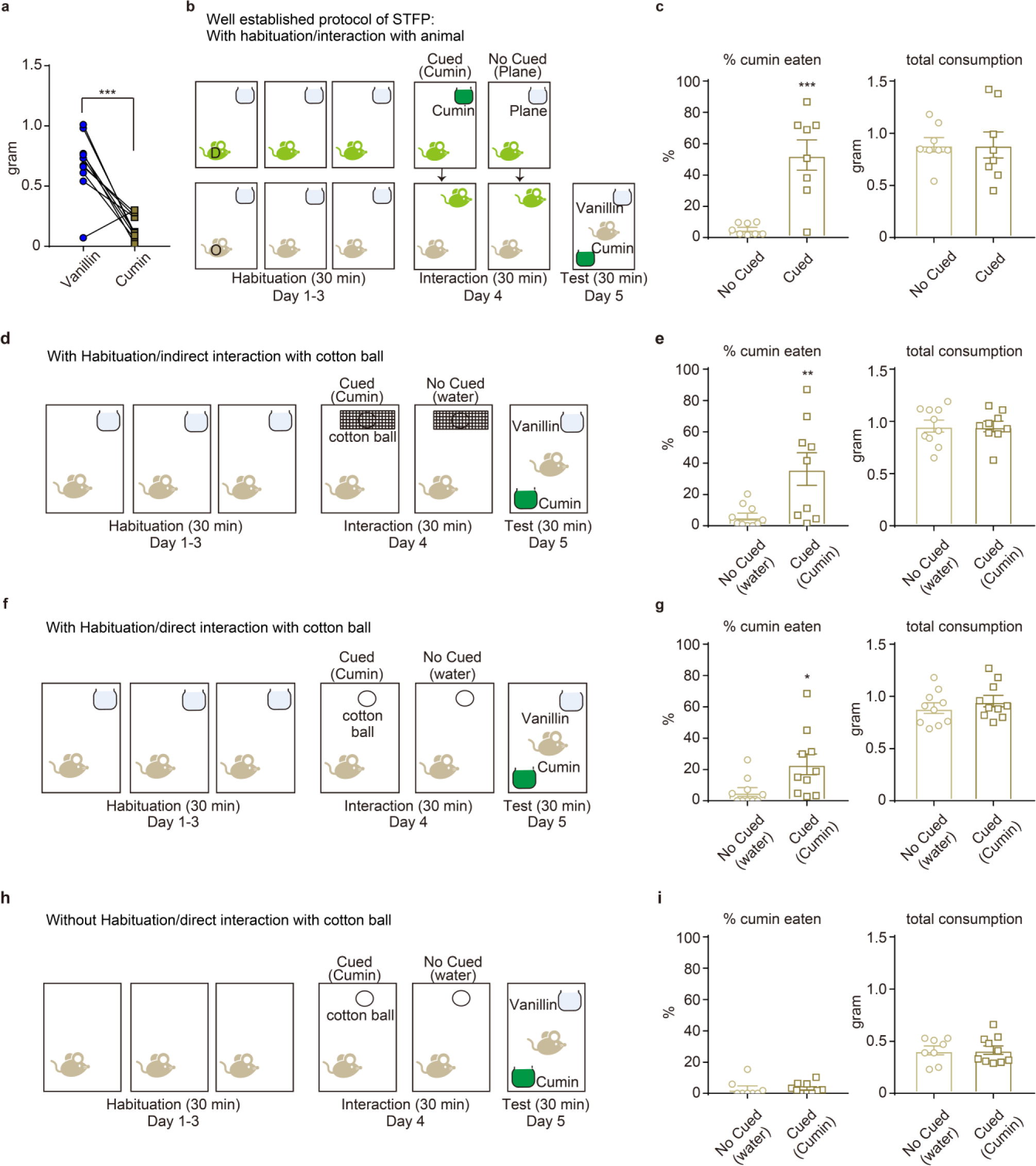
Food preference in the STFP test is driven by non-social effects. **(a)** Natural preference between 0.2% vanillin-odorized and 0.5% cumin-odorized food in mice (n = 10, each group. Two-tailed unpaired *t*-test, t_9_ = 4.958, ***P = 0.0008). **(b)** Schematic illustration of the protocol of STFP which have been well established. D, demonstrator; O, observer. **(c)** Results during the test session. n = 8, each group. % cumin eaten (Left; two-tailed unpaired *t*-test, t_14_ = 4.845, ***P = 0.0003) and total consumption of the odorized food in the jar (Right; two-tailed unpaired *t*-test t_14_ = 0.01753, P = 0.9863). **(d)** Schematic illustration of the protocol. In this case, observer mice interacted with cumin odorized cotton ball instead of the demonstrator during the interaction session. D, demonstrator; O, observer. **(e)** Results during the test session. No Cued, n = 10; Cued, n = 9. % cumin eaten (Left; two-tailed unpaired *t*-test, t_17_ = 2.985, **P = 0.0083) and total consumption of the odorized food in the jar (Right; two-tailed unpaired *t*-test, t_17_ = 0.06343, P = 0.9502). **(f)** Schematic illustration of the protocol. In this case, the odorized cotton ball was surrounded with net to avoid mice from directly touching cumin powder on the cotton. **(g)** Results during the test session. No Cued, n = 10; Cued, n = 9. % cumin eaten (Left; two-tailed unpaired *t*-test, t_18_ = 2.463, *P = 0.0241) and total consumption (Right; two-tailed unpaired *t*-test, t_18_ = 0.8994, P = 0.3803). **(h)** Schematic illustration of the protocol. In this case, mice did not habituate with the jar during the habituation session. **(i)** Results during the test session. No Cued, n = 8; Cued, n = 10. % cumin eaten (Left; two-tailed unpaired *t*-test, t_16_ = 0.05924, P = 0.9535) and total consumption (Right; two-tailed unpaired *t*-test, t_16_ = 0.1764, P = 0.8622).

### Establishment of a novel social learning test - Social Transmission of Food Finding test

We therefore sought to design an improved social learning test that required direct interaction with a demonstrator mouse, named social transmission of food finding (STFF) (Fig. 3). In the STFF, the habituation session with the jar and plain food was omitted for the observer (Fig. 3a) to avoid any prior knowledge about the novel odor and food through firsthand experience-dependent learning (Fig. 1 and 2). Twenty-four hours following the last day of the habituation session (novel powder food in a jar for demonstrator, no jar for observer), the demonstrator mice were exposed to the odorized food-(vanillin or cumin) filled jar for 30 min, for four consecutive days. The consumption of the odorized powder food during this exposure was comparable between the vanillin and cumin groups of demonstrator mice (Fig. 3b). Immediately after this, the demonstrators were placed in the observer mice’s cage for 30 min (interaction session). The time spent nearby the demonstrator and the observer mice during the interaction session was comparable between the odor groups (vanillin and cumin) and the no odor group (Fig. 3c), suggesting that the presence of the odor and the difference of the odors did not influence the duration of social interaction between the demonstrator and the observer mice. Then, during the test session, the observer mice were exposed the vanillin-odorized powder food-filled jar for 30 min. The vanillin group of mice showed a significantly faster latency to start eating and larger consumption of the food in the jar compared to the other groups (Fig. 3d-e). These data suggested that the vanillin group of mice learned that the vanillin odor emitting from the demonstrator mice was food-related during the interaction session.

**Fig. 3:**
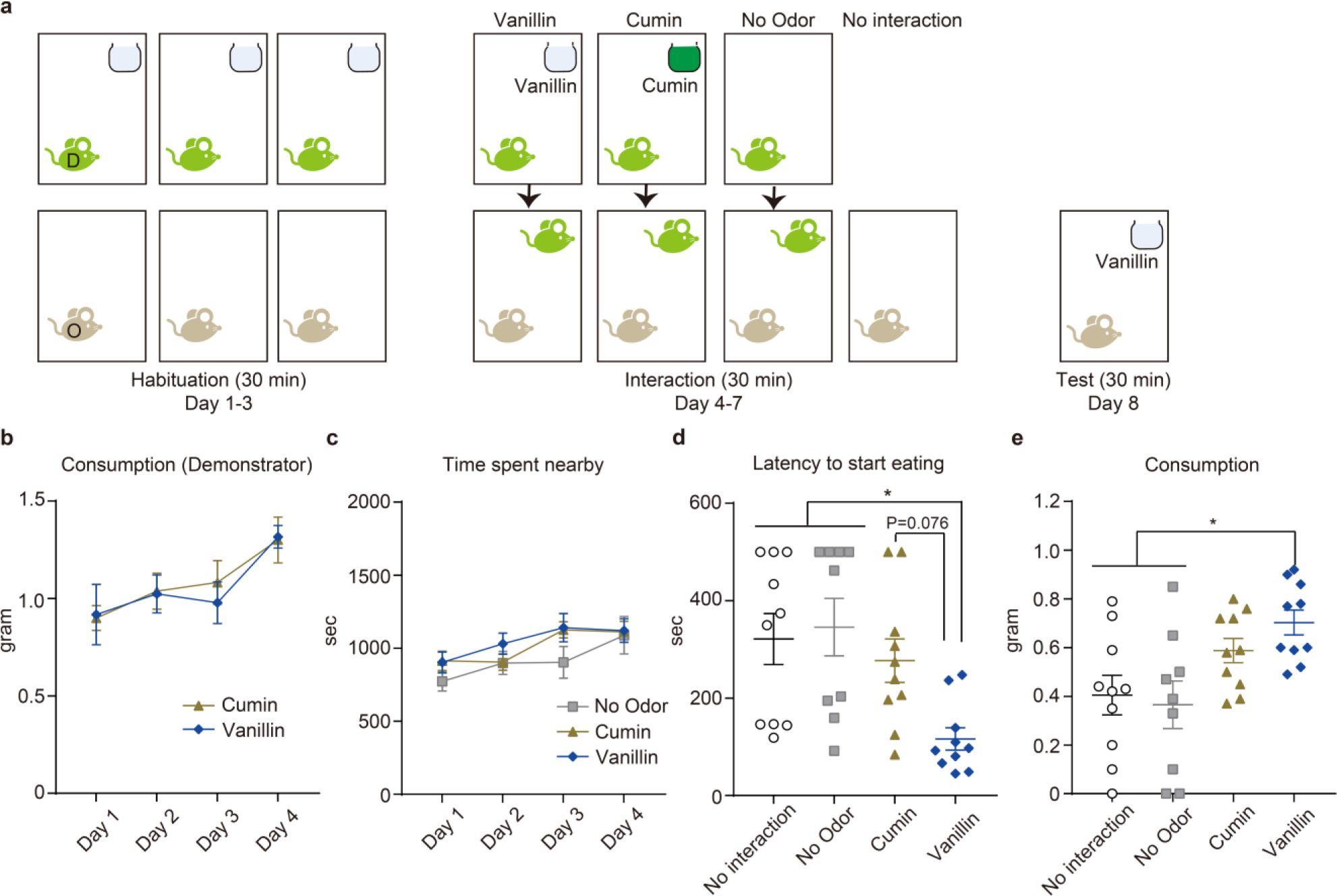
Establishment of a novel social learning test - Social Transmission of Food Finding (STFF) test. **(a)** Schematic illustration of the protocol. D, demonstrator; O, observer. **(b)** Daily consumption of the food by demonstrator mice before interaction with observer mice during the interaction session (No interaction, n = 10; No Odor, n = 9; Cumin, n = 10; Vanillin, n = 10. Two-way repeated ANOVA: Group F_1,72_ = 0.08099, P = 0.7768; Session F_3,72_ = 5.27, **P = 0.0024; Interaction F_3,72_ = 0.1504, P = 0.9291). **(c)** Daily time spent nearby between mice during the interaction (Two-way repeated ANOVA: Group F_2,104_ = 2.814, P = 0.0646; Session F_3,104_ = 5.511, **P = 0.0015; Interaction F_6,104_ = 0.5332, P = 0.7819). **(d)** Latency to start eating the food in the jar during the test session for observer mice (One-way ANOVA; F_3,35_ = 5.068, **P = 0.0051: *post-hoc* Turkey-Kramer test; No mouse vs Vanillin *P = 0.0149, No Odor vs Cumin **P = 0.0071). **(e)** Consumption of the food in the jar for observer mice during the test session (One-way ANOVA: F_3,35_ = 4.9, **P = 0.0060: *post-hoc* Turkey-Kramer test; No mouse vs Vanillin ‘P = 0.0247, No Odor vs Vanillin *P = 0.0114).

### Testing the requirement for social interaction in STFF

In the STFP test, even when the observer mice were exposed to a cumin-odorized cotton ball instead of the demonstrator mice during the interaction session, mice exhibited a preference for cumin-odorized powder food compared to the control no odor group that was either indirectly (Fig. 2d-e) or directly (Fig. 2f-g) exposed to the cotton ball. It has also been previously shown that carbon disulfide (CS_2_), which is one of the odor chemicals from the stomach in rodents, plays a critical role in showing a preference for the odorized food in the STFP test^12,13^. Therefore, we investigated whether an odor itself is enough to establish STFF. Mice were exposed to the cotton ball for four consecutive interaction sessions (Fig. 4a). The cotton balls were odorized with water (No odor), vanillin, CS_2_ or vanillin/CS_2_. The data showed that there was a comparable latency to start eating and consumption of the food in the jar during the test session between groups (Fig. 4b). These results suggest that the memory acquired in the STFF test, measured by the latency to start eating the food and the volume of food consumption, requires direct interaction with the demonstrator.

**Fig. 4:**
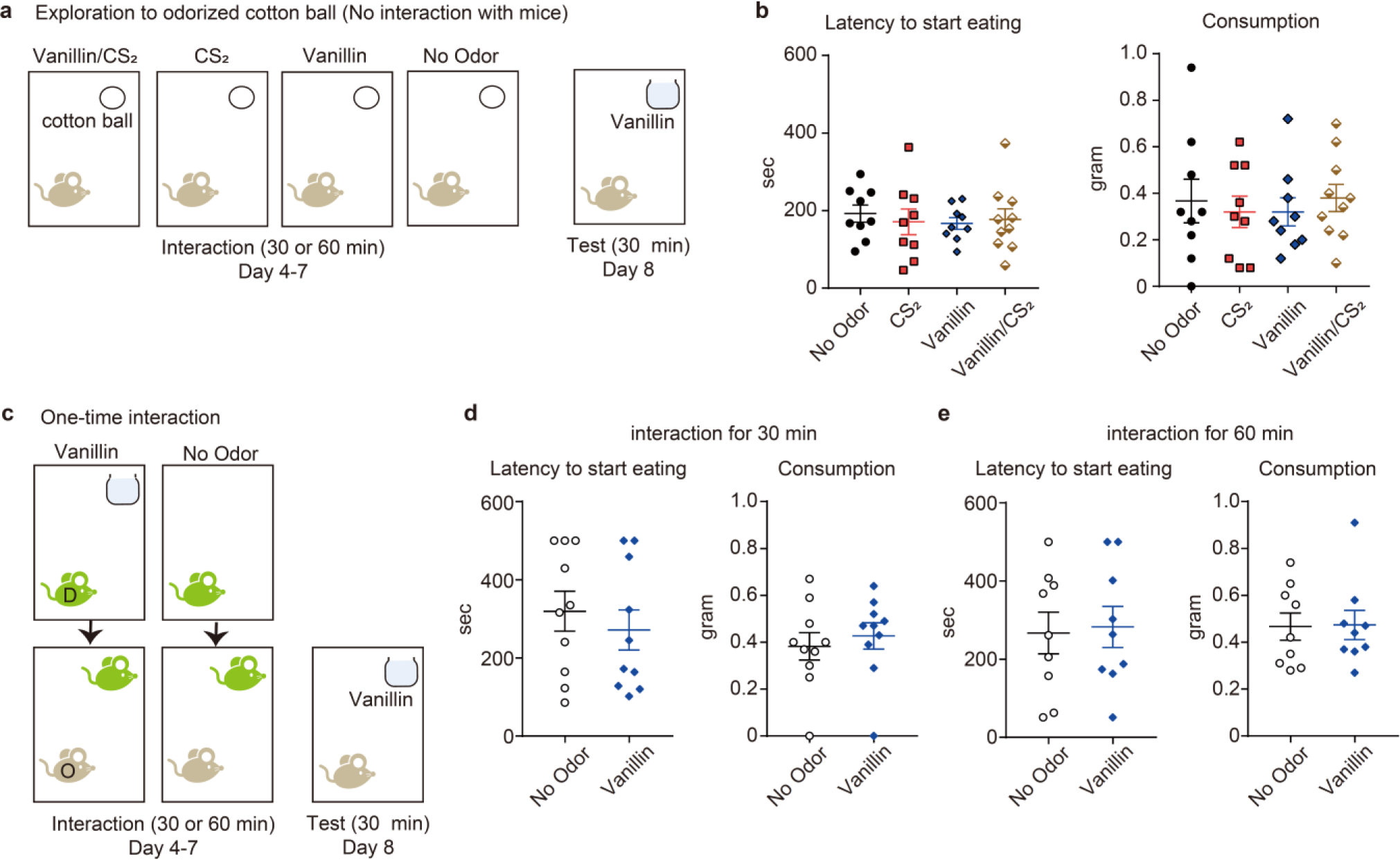
Testing the requirement for social interaction in the STFF. **(a-b)** Analysis the effect of interaction with cotton ball instead of demonstrator mice during the interaction session on feeding behavior. **(a)** Schematic illustration of the protocol. **(b)** Results of STFF. No Odor, n = 9; CS_2_, n = 9; Vanillin, n = 9; Vanillin/CS_2_, n = 10. Latency to start eating (Left: one-way ANOVA; F_3,33_ = 0.1859, P = 0.9052) and consumption of the food in the jar (Right: one-way ANOVA; F_3,33_ = 0.1969, P = 0.8977) during the test session. **(c-e)** Analysis the effect of one interaction on feeding behavior. **(c)** Schematic illustration of the protocol. D, demonstrator; O, observer. **(d-e)** Results of STFF when observer mice interacted with demonstrator mice for 30 min (d) or 60 min (e) during the interaction session. No Odor (30 min), n = 10; Vanillin (30 min), n = 10; No Odor (60 min), n = 9; Vanillin (60 min), n = 9. Latency to start eating (Left: two-tailed unpaired *t*-test; 30 min, t_18_ = 0.6643, P = 0.5149; 60 min, t_16_ = 0.2092, P = 0.8369) and consumption of the food in the jar (Right: two-tailed unpaired *t*-test; 30 min, t_18_ = 0.555, P = 0.5858; 60 min, t_16_ = 0.00786, P = 0.9383) during the test session.

To investigate whether a single interaction with the demonstrator is sufficient to establish the STFF, the observer mice were interacted with the demonstrator only one time, either for 30 min or 60 min long (Fig. 4c-e). Whether for a short (30 min) or long (60 min) interaction, both the latency to start eating and consumption of the food in the jar during the test session were comparable between groups, suggesting that a single interaction was not enough to acquire novel knowledge without firsthand experience in mice, in contrast to generating the food preference in the STFP test.

### STFF test depends on the demonstrator’s context

Previous studies showed that even when observer mice interact with a deceased or physically distressed demonstrator (administered with lithium chloride) they still display a preference for the odor that that the demonstrator had previously consumed in the STFP test^19,20^, indicating that food preference in the STFP test is unaffected by the demonstrator’s health condition or context. Next, we let the observer mice interact with sick demonstrator mice during the interaction session in the STFF (Fig. 5a). To simulate the sick demonstrator mice, they were administered with pentylenetetrazole (PTZ) immediately before interaction with the observer to induce tonic seizure in front of the observer^21^. During the test session, the group of mice which interacted with saline-injected demonstrator mice (saline group) showed a significantly faster latency to start eating the food in the jar compared to the no odor group that the observer interacted with the demonstrator, in which had not consumed any foods before interaction, consistent with the results described above (Fig. 5b left). The latency to start eating in the group of mice interacted with the sick demonstrator (PTZ group) was similar to the no odor group and significantly faster compared to the saline group. However, interestingly, the volume of food consumption in the jar in PTZ group was significantly larger compared to the no odor group and similar to the saline group (Fig. 5b right). These data suggested that although PTZ group of mice recognized vanillin odor was food-related, they hesitated to eat vanillin-odorized food in the jar. Together, STFF test demonstrates that mice can observationally learn positive and negative valence about an unknown odor, depending on the demonstrator’s state, even when they have no firsthand experience with it.

**Fig. 5:**
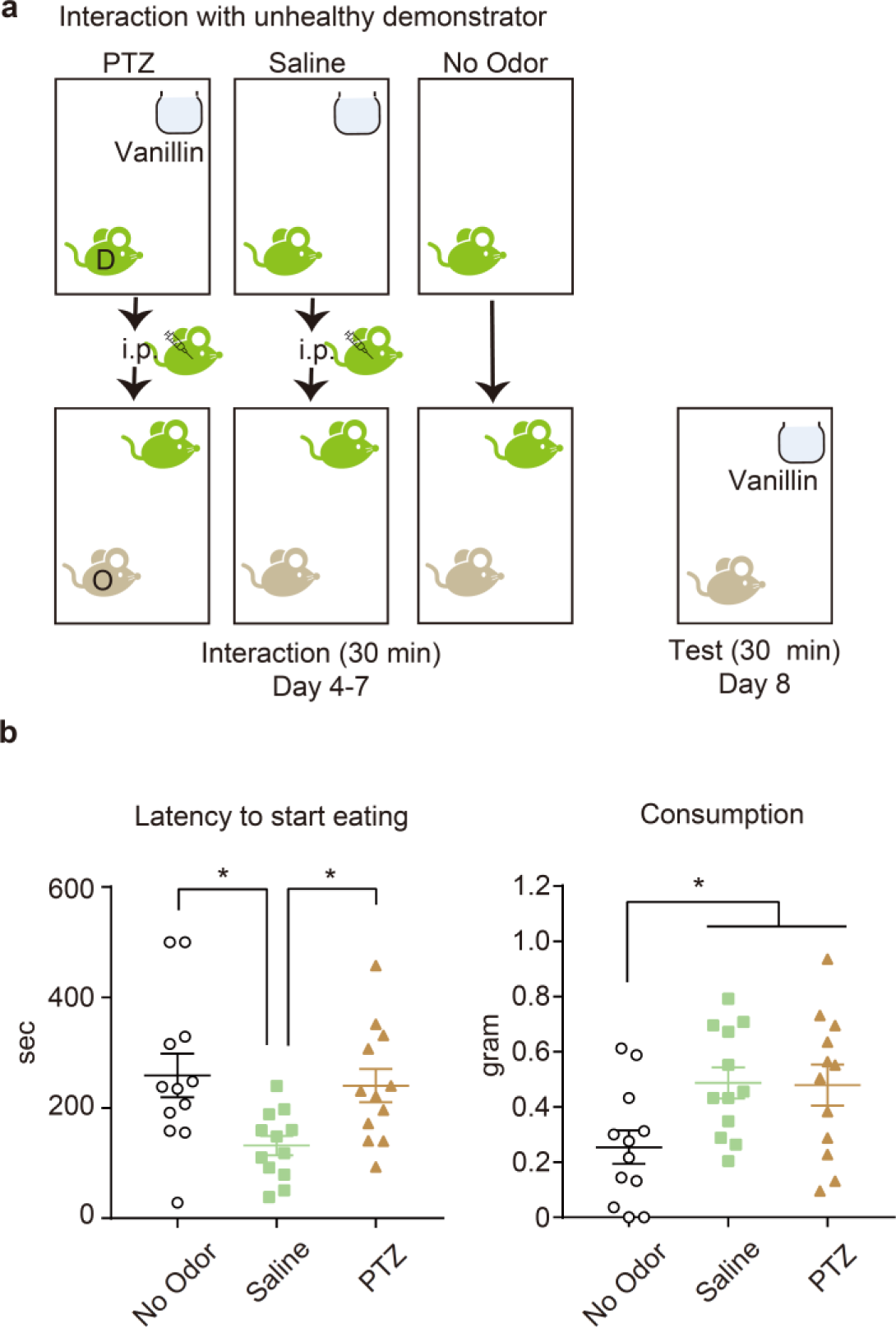
The STFF test depends on the demonstrator’s context. **(a)** Schematic illustration of the protocol. D, demonstrator; O, observer. **(b)** Results of STFF. n = 12, each group. Latency to start eating (Left: one-way ANOVA, F_2,33_ = 5.048, *P = 0.0122; *post-hoc* Turkey-Kramer test, No Odor vs Saline *P = 0.0160, Saline vs PTZ *P = 0.0439) and consumption of the food in the jar (Right: one-way ANOVA, F_2,33_ = 4.236, *P = 0.0230; *post-hoc* Turkey-Kramer test, No Odor vs Saline *P = 0.0391, No Odor vs PTZ *P = 0.0476) during the test session.

### Roles of dorsal hippocampal CA1 in STFF

To investigate the role of the dorsal hippocampus in the formation of STFF memory, the dorsal hippocampus of the observer was chemically or genetically lesioned before the interaction sessions (Fig. 6a-b). The microinjection of ibotenic acid (IBO)^22^ into the dorsal hippocampal CA1 region (dCA1) resulted in a global lesion of the dorsal hippocampus including CA1, CA2, CA3 and dentate gyrus (Fig. 6c). Under this condition, the cued group of mice that interacted with the demonstrator that had consumed the vanillin odorized powder food before interaction exhibited a comparable latency to start eating and consumption of the food in the jar during the test session compared to the Non-cued group of mice which interacted with the demonstrator that had eaten no food before interaction, despite the sham (vehicle injected)-cued group of mice showed a significantly faster latency to start eating and a significantly larger consumption of the food (Fig. 6d). To further examine the roles of the dCA1 on STFF, we injected a cocktail of Cre expressing and Cre-dependent tacaspase3 expressing AAVs^23^ into the dCA1 of the observer, which caused dCA1-specific lesion (Fig. 6e). Even under this condition, the dCA1-lesioned observer mice did not show faster latency to start eating and larger consumption of the food in the jar during the test session (Fig. 6f). These data suggested that the dCA1 of the hippocampus is necessary for formation of STFF memory.

**Fig. 6:**
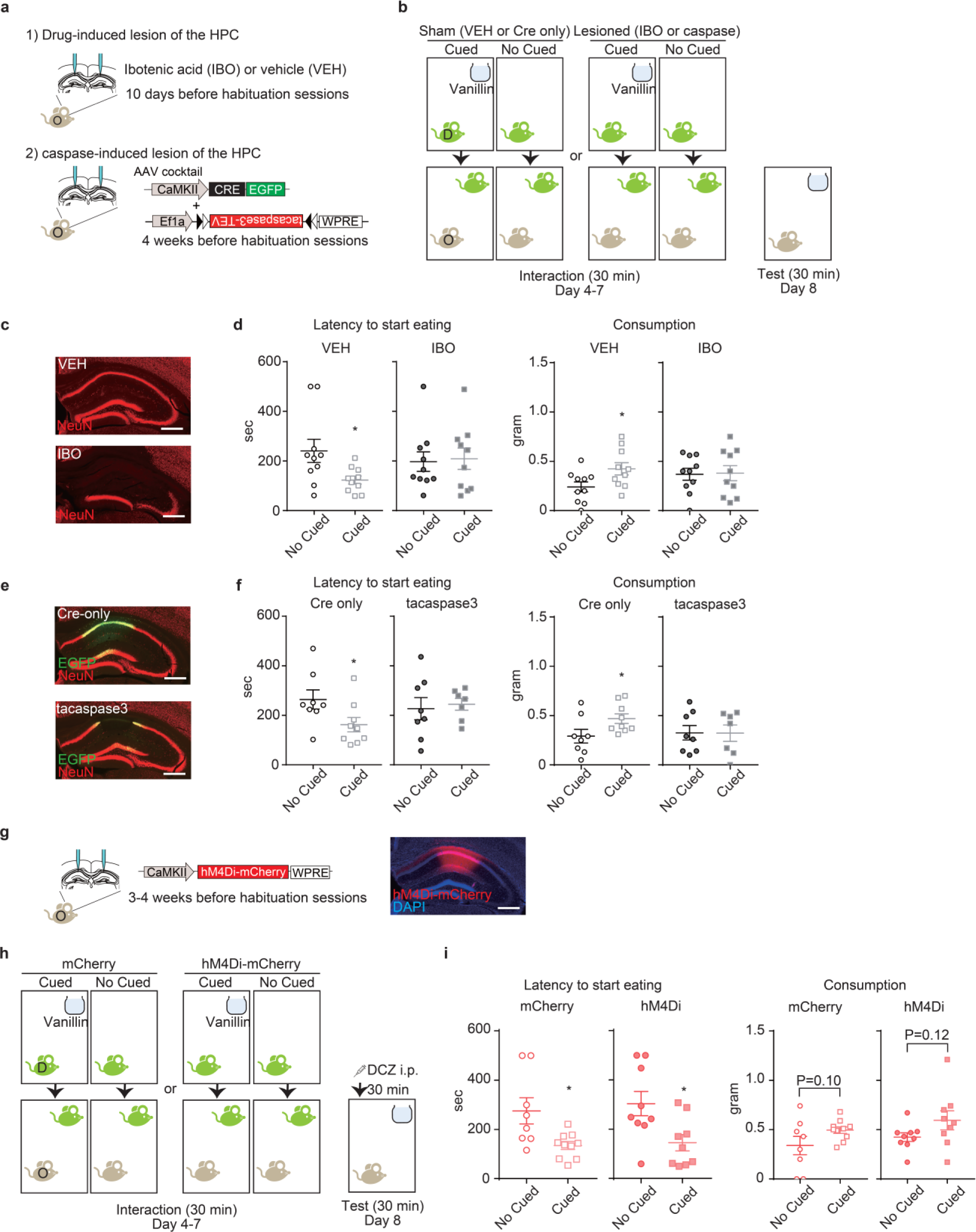
Roles of the dorsal hippocampal CA1 in the STFF. **(a-f)** Analysis the role of the dorsal hippocampal CA1 (dCA1) in the formation of STFF memory. **(a)** There used two methods of the lesion of the dCA1, injection of ibotenic acid (IBO) (upper)-, or AAV-inducible expression of caspase (lower)-dependent lesion. **(b)** Schematic illustration of the protocol. D, demonstrator; O, observer. **(c)** The effect of IBO injection into the dCA1. Successful lesion of CA region including the dentate gyrus of the hippocampus using IBO. The brain sections were stained with anti-NeuN (red). **(d)** Results of STFF. VEH, vehicle. VEH-No Cued, n = 10; VEH-Cued, n = 11; IBO-No Cued, n = 10; IBO-Cued, n = 10. The latency to start eating (Left: two-tailed unpaired *t*-test; VEH, t_18_ = 2,374, *P = 0.0289, IBO, t_18_ = 0.2033, P = 0.8412) and the consumption of odorized powder food in the jar (Right: two-tailed unpaired *t*-test; VEH, t_18_ = 2.312, *P = 0.0328, IBO, t_18_ = 0.1134, P = 0.9109) during the test session. **(e)** The effect of AAV-cocktail injection into the dCA1. The brain sections were stained with anti-NeuN (red) and anti-GFP (green). **(f)** Results of STFF. Cre only-No Cued, n = 8; Cre only-Cued, n = 9; tacaspase3-No Cued, n = 8; tacaspase3-Cued, n = 7. The latency to start eating (Left: two-tailed unpaired *t*-test; Cre only, t_15_ = 2.139, *P = 0.0493, tacaspase3, t_13_ = 0.3336, P = 0.7440) and the consumption of odorized powder food in the jar (Right: two-tailed unpaired *t*-test; Cre only, t_15_ = 2.132, *P = 0.0500, tacaspase3, t_13_ = 0.03253, P = 0.9745) during the test session. **(g-i)** Analysis the role of the dCA1 in retrieval of STFF memory. **(g)** hM4Di-mCherry expressing AAV was injected with dCA1. The brain sections were stained with anti-mCherry (red) and DAPI (blue) after STFF. **(h)** Schematic illustration of the protocol. D, demonstrator; O, observer. The effect of the inhibition of dCA1 using G(i)-DREADD during retrieval of STFF. **(i)** Results of STFF. mCherry-No Cued, n = 8; mCherry-Cued, n = 10; hM4Di-No Cued, n = 9; hM4Di-Cued, n = 9. The latency to start eating (Left: two-tailed unpaired *t*-test; mCherry, t_16_ = 2.796, *P = 0.0129, hM4Di, t_16_ = 2.658, *P = 0.0172) and the consumption of odorized powder food in the jar (Right: two-tailed unpaired *t*-test; mCherry, t_16_ = 1.723, P = 0.1042, hM4Di, t_16_ = 1.606, P = 0.1279) during the test session. Scale bars in the images are 500 µm.

In keeping with previous studies^24–26^, we confirmed that the hippocampus was necessary for both the formation and retrieval of STFP memory, since the activation of hM4Di DREADD^27^ in the dCA1 of the hippocampus with DCZ, an agonist of the muscarine receptor DREADD^28^, during the interaction session or the test session reduced the preference in the STFP test (Supplemental Fig. 1a-e). However, interestingly, dCA1 was dispensable for the recall of the odor itself-dependent preference (Supplemental Fig. 1f-g). Taken together, sniffing the odor from the demonstrator is a critical step required for dCA1-dependent socially-transmitted memory formation.

Finally, we investigated the role of the dCA1 in the retrieval of STFF memory. The hM4Di-, or mCherry-expressing AAV was injected into the dCA1 of the observer animals 3-4 weeks before the habituation session (Fig. 6g). Thirty-minutes before the test session, the observer was administered with DCZ under anesthesia to exclude any stress caused by its injection (Fig. 6h). Both the mCherry-Cued and hM4Di-Cued groups of mice showed a significantly faster latency to start eating and larger consumption of the food in the jar during the test session (Fig. 6i), suggesting that the dCA1 of the hippocampus was dispensable for the retrieval of STFF memory. Altogether, dorsal hippocampal CA1 region is differently involved during the learning and retrieval processes for STFF.

## Discussion

In this study, we first investigated the critical factors for the classic STFP test (Fig. 2). We found that the odor itself (without the demonstrator) was sufficient to generate a preference for the odor in the STFP test. Furthermore, the preference generated from the odor itself was not observed when the observer had no habituation session with the food-filled jar, which has employed in the protocol of STFP^9,16,17^. These findings indicate that the food preference in the STFP test can be driven by non-social components, such as familiarity with the odor itself and prior direct experience with a food-filled cup. These findings also suggest that the co-incidence of jar-food memory and familiarity with the unknown odor in the test session might induce a preconception of odor-food association or generate a false associative memory. Second, we examined the role of dCA1 neurons in the STFP test (Supplemental Fig. 1). We found that the preference generated by interaction with the demonstrator was dCA1- dependent; however, the preference formed by sniffing the odor alone (without the demonstrator) was not dCA1-dependent. The olfactory bulb directly innervates the entorhinal cortex, which cooperates with dCA1 under odor stimuli^29,30^. We believe that the dCA1 integrates the episodes, including the experience of exploring the food in the jar, the experience of smelling the odor from the other, and the unknown odor, to generate a preference in the STFP^31,32^. Together we conclude that the food preference in the STFP test is facilitated by both the socially-induced factor (dCA1-dependent) and firsthand experience-induced factor (dCA1-independent).

Next, we designed an improved social learning task, named social transmission of food preference (STFF), by modifying the STFP test. The STFF allowed the observer to acquire new knowledge of which the novel odor is food-related from other animals, even when the observer has no prior knowledge about the odor and food in their life (Fig. 3). In contrast to the STFP test, the STFF required multiple social interactions to obtain STFF memory (Fig. 4). Importantly, we revealed that mice could recognize the demonstrator’s context in the STFF test. We found that interacting with the animal showing PTZ-induced tonic seizure or dying impaired induced faster latency to start eating, proving that mice understand the meaning and valence of the odor based on the other animal’s situation, healthy or sick. However, once the observer eats the food and realizes it is safe based on their own experience during the test session, they showed increased consumption, suggesting that the observer has knowledge that the odor is food-related (Fig. 5). Understanding the other’s context and behavior through observation is vital for animals to survive in nature. Future studies must investigate how animals understand the positive or negative implications conveyed by the other animals without firsthand experience. These data conclude that the STFF is suitable for assessing valence-rich social learning in mice and can be helpful in examining how the observer socially recognizes the demonstrator’s condition and context.

Previous studies have shown that dorsal hippocampal CA2 region (dCA2), including the lateral entorhinal cortical-dCA2-ventral CA1 circuit, governs social information such as social recognition and social novelty^33–35^. Our current study showed that dCA1 is required for acquiring new knowledge by socially interacting with the demonstrator in STFF test (Fig. 6). Additionally, we found that establishing STFF requires several interactions with the demonstrator (Fig. 4). These findings suggest that the STFF is selected based on the observer’s self-experience of smelling the unknown but the same odor from the other animals rather than social information such as the identity of the demonstrator. However, we do not deny the possibility that social information also contributes to STFF through other regions, such as dCA2 and ventral CA1^36^.

How does the observer animal understand that the unknown odor is food-related? We tested the effect of carbon disulfide (CS_2_), one of the odor chemicals from the rodents’ stomachs, plays a critical role in recognizing odor as food-related without the demonstrator. However, we did not observe any effect on the latency to start eating (Fig. 4). Therefore, we currently speculate that mice may use memory schemas about food intake previously acquired. Schemas are cognitive structures that systematically organize and understand the knowledge that animals have experienced and developed throughout their lives and influence new memory formation^37^. Thus, mice may store vast information about food intake as a schema from birth^38^. During the interaction with the demonstrator, the observer animal may realize that the odor is food-related by activating the schemas network, including the medial prefrontal cortex^39,40^. Future studies must explore this possibility by devising ways to block the formation of food-intake schemas, such as by preventing mice from group housing since birth.

Our study showed that the lesion of dCA1 before the interaction session, but not inhibition of dCA1 activity during the test session, impaired the induction of faster latency to start eating the food in the jar, suggesting that the responsible region for STFF memory rapidly changes from the hippocampus to another area, such as the para-hippocampal or cortical region (Fig. 6). Why does this happen? First, we consider that the way of recalling the odor differs between the interaction and test sessions. Previous studies suggest that episodic memories are recalled from different perspectives in humans; own-eyes (egocentric) or observer perspective^41^. It has been suggested that the own-eyes perspective is more vivid than the observer perspective and that the regions involved in the recall of each perspective may be different^42,43^. During the interaction session, the observer animal may repeatedly recall their experience of interacting with other animals and the unknown but the same odor from their own-eyes perspective. On the other hand, the memory of the unknown odor is recalled from their observer perspective rather than their own-eyes perspective during the test session. Second, we consider that STFF memory is transformed from episodic-like to semantic-like^44^. Some evidence has showed that different regions are involved in episodic and semantic memories^45,46^ (but also see^47^). Thus, similar information may be extracted from multiple experiences and stored as semantic memory-like memories in regions other than the hippocampus. We showed that multiple (four times), but not a single, interaction with the demonstrator was required for the establishment of STFF (Fig. 3). We speculate that the formation of STFF memory may require a systems consolidation process^48,49^ through multiple interaction episodes with the demonstrator, which allows the observer to obtain new knowledge from the demonstrator. Further studies need to investigate these possibilities by identifying the brain regions involved in the recall of STFF memory and measuring the brain activities in the such areas including dCA1, para-hippocampal and cortical regions during STFF.

## Methods

### Animals

All procedures relating to mouse care and experimental treatments conformed to NIH and Institutional guidelines and were conducted with the approval of the UT Southwestern Institutional Animal Care and Use Committee (IACUC). All experiments used wild-type male mice of the C57BL/6J at least over 8 weeks old. Mice were group housed with littermates for a minimum of 7 days before food deprivation in a 12 h (6 am-6 pm) light/dark cycle, with food and water available *ad libitum*. For food deprivation, all mice were housed in a single and were supplied with 2.0 – 2.5 g of food per day at least for 3 days prior to experiments. During experiments, the weights of mice were monitored, and the food was supplied to be 80 % of their original weight. All experiments were conducted during morning. All mice were housed in the same size of cage (30.0 cm W × 16.0 cm H × 14.5 cm H). Mice were randomly assigned to experimental conditions.

### Social Transmission of Food Preference (STFP)

Mice were housed in groups of four. After food deprivation as described above, mice were separated into the demonstrator or the test observer groups two by two per cage, and all mice were single housed. During the habituation session, the demonstrator group and the observer group of mice were entered into a new cage with one or two novel powder food (AIN-93M, Bio-Serv) -filled feeding jars (Dyets Inc.), respectively, for 30 min for 3 consecutive days (the habituation session). Twenty-four hours following, the demonstrator was entered into a new cage with 0.5 % cumin (McCormick)-containing powder food-filled jar for 30 min, and immediately after this procedure (within 1 min), the demonstrators were then entered into the observer’s cage for 30 min (the interaction session). Since mice naturally preferred for vanillin-odorized than cumin-odorized food (Fig. 2a), the demonstrators were subjected to cumin-odorized food during the interaction session as recommended in previous report ^9^. All demonstrator animals were cage mates for observer to avoid severe fighting. The test session was performed 24 hrs following the interaction session. The observer animals were entered into a new cage with both cumin- and vanillin (0.2 %, Sigma-Aldrich)-containing powder food-filled jar for 30 min. Consumption of these powder food during the test session was measured compared to the original amount. The percent cumin eaten in figures were calculated as following:

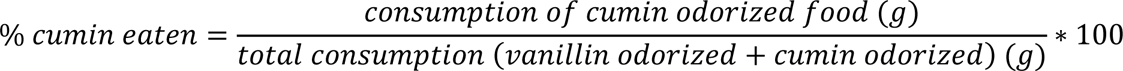

For the non-interaction with the demonstrator experiment (Fig. 2e-j), the odorized cotton ball instead of the demonstrator animals were provided to the observer animal during interaction session. Immediately after the interaction session, the cotton balls were removed from the cage, and the cage was changed to a new one. To make odorized cotton ball, cotton balls were dipped into 0.5 % cumin -containing water or just water, and then were completely dried under hood before STFP. The net was used for surrounding the balls to prevent mice from touching cotton balls directly.

For neuronal inhibition experiments using hM4D(Gi)-DREADD (Supplemental Fig. 1), mice were injected with AAV_2/8_-CaMKII-hM4D(Gi)-mCherry (Addgene, titer: 2.3*10^13^ genome/mL) or AAV_2/5_-CaMKII-mCherry (UNC Vector Core, titer: 2.3*10^12^ genome/mL) into the dorsal part of CA1 (dCA1) of hippocampus. Three to four weeks following this, mice were subjected to STFP task. Mice were administrated with DCZ (100 µg/kg, i.p.) 30 min before the interaction or the test session to analyze the role of dCA1 in formation or retrieval of STFP memory, respectively, under anesthetizing with isoflurane (1.0 - 1.5 %).

The animal behaviors were video monitored during the interaction and the test session. The time spent nearby between the animals during the interaction session and the time spent on the jar during the test session were analyzed using UMA Tracker^50^ (data not shown). There were no statistically significant about weight before test session between groups in all experiments (Supplemental table 1). Following checking the expression by immunohistochemistry, we excluded the sample from the data which the AAV solution was leaked out other than dCA1 region in Supplemental Fig. 1. All mice were blinded by experimenter during the test session and analyzing the video data.

### Social Transmission of Food Finding (STFF)

Mice were housed in groups of four, except the experiment of interaction with sick demonstrator (Fig. 5, described below). After food deprivation as described above, mice were separated into the demonstrator or the test observer groups two by two per cage, and all mice were single housed. During the habituation session, the demonstrator animals were subjected the same as in the STFP described above, while the observer animals were entered an empty new cage for 30 min for 3 consecutive days. The interaction session was performed the same as STFP, but for 4 consecutive days for 30 min per day (Fig. 3g-k), or for 1 day for 30 min or 60 min (Fig. 4c-e). The demonstrator animals were changed daily so that the observer animals interacted twice per one demonstrator animal through 4 consecutive interaction sessions. During the test session, the observer animals were entered into a new cage with one vanillin-odorized powder food-containing feeding jar for 30 min. The latency to start eating the powder food in the jar and consumption of the food were measured. The latency to start eating was defined as mice ate the powder food at the first. This was clearly defined since almost mice showed a “chewing” behavior (Supplemental video 1). The latency to start eating was set to a maximum of 500 seconds.

For the non-interaction with the demonstrator experiment (Fig. 4a-b), the odorized cotton ball was entered into the observer’s cage for 4 consecutive days for 30 min during the interaction session. Immediately after the interaction session, the cotton ball was removed, and the cage was changed to a new one. To make the odorized cotton balls, the cotton balls were dipped into 0.2 % vanillin/13 µM CS_2_-, vanillin-, CS_2_-containing water, or just water, and then were completely dried under hood before STFF.

For the experiment of interaction with unhealthy demonstrator animals (Fig. 5), mice were housed in groups of five. After food deprivation, four of them were used as the demonstrator and the other was used as the observer. All mice were single housed. The PTZ group of demonstrator animals was administrated with pentylenetetrazol (PTZ, Sigma-Aldrich) before the interaction session (30 mg/kg, i.p. and 10 mg/kg, i.p. at 2 days and at 1 day before the interaction session, respectively) to easily induce tonic seizure. Immediately after eating the powder food during the interaction session, the demonstrator was administrated with high dose of PTZ (60 mg/kg, i.p.), and then was entered into the observer’s cage for 30 min. Around 5 min after entering, PTZ-injected demonstrator animals showed the tonic seizure, and some became a dead during the interaction session. The demonstrator animals were changed daily so that the observer animals interacted different demonstrator animal through 4 consecutive interaction sessions in this experiment.

For the experiment of dCA1 lesion to investigate the role of dCA1 in formation of STFF (Fig. 6a-f), the observer animal were injected with ibotenic acid (IBO, 10 mg/mL) or AAV-cocktail of AAV_2/8_-CaMKII-Cre-EGFP (UNC Vector Core, final titer: 2.15*10^12^ genome/mL) and AAV_2/1_-Ef1a-flex-taCasp3-TEVp (UNC Vector Core, final titer: 1.6*10^12^ genome/mL) into the dCA1 of hippocampus 10 days or 4 weeks before the habituation session, respectively. As the control, the vehicle solution for IBO or AAV_2/8_-Cre (titer: 2.15*10^12^ genome/mL) for AAV-cocktail were used.

For the experiment to investigate the role of dCA1 in retrieval of STFF (Fig. 6g-i), the observer animals were injected with AAV_2/8_-CaMKII-hM4D(Gi)-mCherry or AAV_2/5_-CaMKII-mCherry into the dCA1 of hippocampus. Three to four weeks following this, mice were subjected to STFF task. Mice were administrated with DCZ (100 µg/kg, i.p.) 30 min before the test session under anesthetizing with isoflurane (1.0 - 1.5 %).

The animal behaviors were video monitored during the interaction and the test session. The time spent nearby between the animals during the interaction session was analyzed using UMA Tracker^50^. There were no statistically significant about weight before the test session between groups in all experiments (Supplemental table 1). Following checking the expression by immunohistochemistry, we excluded the sample from the data which the AAV solution was leaked out other than dCA1 region in Fig. 6. All mice were blinded by experimenter during the test session and analyzing the video data.

### Drugs

All drugs were stored at -20 ℃ until use. Deschloroclozapine (DCZ, hellobio) was dissolved at 10 µg/mL in PBS. Ibotenic acid (IBO, Sigma-Aldrich) was dissolved at 10 mg/mL in PBS. PTZ was dissolved at 1, 3 or 6 mg/mL in PBS. DCZ and PTZ were melted at least 1 hr before intraperitoneal (i.p.) injection to be a room temperature (R.T.).

### Stereotaxic Surgery

All surgeries were performed as previously described^5,6^. All surgeries were conducted using asceptic technique and followed NIH and UT Southwestern IACUC guidelines. A digital small animal stereotaxic (David Kopf Instruments) with a stereomicroscope (Leica) was used to perform all stereotaxic procedures. Mice were anesthetized with isoflurane (1.0 - 1.5%). Microinjections were completed with 10 μL Hamilton microsyringe filled with mineral oil and with a glass micropipette (Drummond Scientific Company) filled with mineral oil attached. A microsyringe pump (World Precision Instruments) was used to control injection speed and volume. The micropipette was slowly lowered to the target site and remained for 5 min after injection. Mice were given meloxicam (2 mg/kg) as a post-surgical analgesic, remained on the heating pad until fully recovered from the anesthesia, and were allowed to recover a minimum of three days before returning to group housing with cage mates. To verify target sites, post-mortem histology was performed. The following coordinate was used for injection into the dCA1 of hippocampus: AP: -2.00 mm; ML: ±1.35 mm; DV: -1.27 mm relative the bregma. The volumes of IBO or its vehicle were 220 nL per side, and that of AAVs were 250 nL per side.

### Immunohistochemistry

Immunohistochemistry was performed as previously described^5,6^. Mice were deeply anesthetized with a ketamine (75 mg/kg)/ dexmedetomidine (1 mg/kg) cocktail and transcardially perfused with 4 % paraformaldehyde (PFA) in PBS. Brains were removed and post-fixed in 4 % PFA in PBS at 4 _°_C for 24 h and then sectioned using a vibratome (Leica) with thicknesses of 40 μm. For immunohistochemistry (IHC), tissue sections were incubated in 0.3 % Triton-X/PBS (PBS-T) with 10 % normal goat serum (NGS) for 1 h. Primary antibodies were then added to PBS-T with 10 % NGS and tissue sections were incubated overnight at 4 _°_C. Primary antibodies used for immunostaining were as follows: guinea pig anti-NeuN (1:1000, Millipore Sigma), rabbit anti-GFP (1:2000, Thermo Fisher Scientific) and rat anti-mCherry (1:2000, Invitrogen). After washing with PBS (3 x 5 min), tissue sections were subsequently incubated with AlexaFluor 488, AlexaFluor 546, or AlexaFluor 555 conjugated secondary antibodies (1/500, Thermo Fisher Scientific) in PBS with 10 % NGS for 2 h at R.T. Tissue sections were then incubated in PBS-T with DAPI (1:1000, Thermo Fisher Scientific) for 20 min, and then washed with PBS-T (3 x 10 min) at R.T., and mounted in VECTASHIELD medium (Vector Laboratories) on glass slides. Fluorescence images were taken with a Zeiss AxioImager M2 microscope with Apotome using the 5x objective. Images were processed using the Zen Blue software.

### Statistical Analysis

Statistical analyses of the acquired data were done with Prism 7 (GraphPad). Paired *t*-test (two-sided), unpaired *t*-test (two-sided) and one-way ANOVA with Turkey-Kramer test were used. Error bars shown in the figures represent the Standard Error of the Mean (SEM). All data analyses were performed blinded for experimenter. The null hypothesis was rejected at the P < 0.05 level.

## Acknowledgments

We thank all members of the Kitamura Laboratory for their support and hot discussion. This work was supported by Endowed Scholar Program at University of Texas Southwestern Medical Center and KAKENHI JP20K06849 (to R.K.) and JP 22H00432 (H.B.). R.K. received support from an International Platform for Young Researchers (The University of Tokyo).

## Author Contributions

R.K. and T.K. designed this study. R.K. performed all experiments. R.K. H.B. and T.K contributed to the interpretation of data. R.K. H.B. and T.K. wrote the paper. All authors approved the final manuscript.

## Completing interests

The authors declare no competing interests.

## References

1. Bandura, A. Social Learning Theory of Aggression. Journal of Communication 28, 12– 29 (1978).

2. Behrens, T. E. J., Hunt, L. T. & Rushworth, M. F. S. The Computation of Social Behavior. Science 324, 1160–1164 (2009).

3. Monfils, M. H. & Agee, L. A. Insights from social transmission of information in rodents. Genes Brain Behav 18, e12534 (2019).

4. Gariépy, J.-F. et al. Social learning in humans and other animals. Front Neurosci 8, (2014).

5. Terranova, J. I., Yokose, J., Osanai, H., Ogawa, S. K. & Kitamura, T. Systems consolidation induces multiple memory engrams for a flexible recall strategy in observational fear memory in male mice. Nat Commun 14, 3976 (2023).

6. Terranova, J. I. et al. Hippocampal-amygdala memory circuits govern experience-dependent observational fear. Neuron 110, 1416–1431.e13 (2022).

7. Keum, S. & Shin, H.-S. Neural Basis of Observational Fear Learning: A Potential Model of Affective Empathy. Neuron 104, 78–86 (2019).

8. Huang, Z. et al. Ventromedial prefrontal neurons represent self-states shaped by vicarious fear in male mice. Nat Commun 14, 3458 (2023).

9. Bessières, B., Nicole, O. & Bontempi, B. Assessing recent and remote associative olfactory memory in rats using the social transmission of food preference paradigm. Nat Protoc 12, 1415–1436 (2017).

10. Posadas-Andrews, A. & Roper, T. J. Social transmission of food-preferences in adult rats. Anim Behav 31, 265–271 (1983).

11. Mou, X., Pokhrel, A., Suresh, P. & Ji, D. Observational learning promotes hippocampal remote awake replay toward future reward locations. Neuron 110, 891–902.e7 (2022).

12. Munger, S. D. et al. An Olfactory Subsystem that Detects Carbon Disulfide and Mediates Food-Related Social Learning. Current Biology 20, 1438–1444 (2010).

13. Galef, B. G., Mason, J. R., Preti, G. & Bean, N. J. Carbon disulfide: A semiochemical mediating socially-induced diet choice in rats. Physiol Behav 42, 119–124 (1988).

14. Farrow, C. & Coulthard, H. 11 - Multisensory evaluation and the neophobic food response. in Food Neophobia (ed. Reilly, S.) 219–236 (Woodhead Publishing, 2018). 10.1016/B978-0-08-101931-3.00011-2.

15. Modlinska, K., Stryjek, R. & Pisula, W. Food neophobia in wild and laboratory rats (multi-strain comparison). Behavioural Processes 113, 41–50 (2015).

16. Loureiro, M. et al. Social transmission of food safety depends on synaptic plasticity in the prefrontal cortex. Science 364, 991–995 (2019).

17. Wrenn, C. C. Social Transmission of Food Preference in Mice. Curr Protoc Neurosci 28, 8.5G.1–8.5G.7 (2004).

18. Galef, B. G., Kennett, D. J. & Stein, M. Demonstrator influence on observer diet preference: Effects of simple exposure and the presence of a demonstrator. Anim Learn Behav 13, 25–30 (1985).

19. Galef, B. G. & Stein, M. Demonstrator influence on observer diet preference: Analyses of critical social interactions and olfactory signals. Anim Learn Behav 13, 31–38 (1985).

20. GALEF JR., B. G. Direct and Indirect Behavioral Pathways to the Social Transmission of Food Avoidancea. Ann N Y Acad Sci 443, 203–215 (1985).

21. Shimada, T. & Yamagata, K. Pentylenetetrazole-induced kindling mouse model. Journal of Visualized Experiments 2018, (2018).

22. Isacson, O., Brundin, P., Kelly, P. A. T., Gage, F. H. & Björklund, A. Functional neuronal replacement by grafted striatal neurones in the ibotenic acid-lesioned rat striatum. Nature 311, 458–460 (1984).

23. Yang, C. F. et al. Sexually Dimorphic Neurons in the Ventromedial Hypothalamus Govern Mating in Both Sexes and Aggression in Males. Cell 153, 896–909 (2013).

24. Bunsey, M. & Eichenbaum, H. Selective damage to the hippocampal region blocks long-term retention of a natural and nonspatial stimulus-stimulus association. Hippocampus 5, 546–556 (1995).

25. Ross, R. S. & Eichenbaum, H. Dynamics of Hippocampal and Cortical Activation during Consolidation of a Nonspatial Memory. The Journal of Neuroscience 26, 4852 (2006).

26. Lesburguères, E. et al. Early Tagging of Cortical Networks Is Required for the Formation of Enduring Associative Memory. Science 331, 924–928 (2011).

27. Roth, B. L. DREADDs for Neuroscientists. Neuron 89, 683–694 (2016).

28. Nagai, Y. et al. Deschloroclozapine, a potent and selective chemogenetic actuator enables rapid neuronal and behavioral modulations in mice and monkeys. Nat Neurosci 23, 1157–1167 (2020).

29. Igarashi, K. M., Lu, L., Colgin, L. L., Moser, M.-B. & Moser, E. I. Coordination of entorhinal–hippocampal ensemble activity during associative learning. Nature 510, 143–147 (2014).

30. Nagayama, S. et al. Differential Axonal Projection of Mitral and Tufted Cells in the Mouse Main Olfactory System. Front Neural Circuits 4, (2010).

31. Dimsdale-Zucker, H. R., Ritchey, M., Ekstrom, A. D., Yonelinas, A. P. & Ranganath, C. CA1 and CA3 differentially support spontaneous retrieval of episodic contexts within human hippocampal subfields. Nat Commun 9, 294 (2018).

32. Schlichting, M. L., Zeithamova, D. & Preston, A. R. CA1 subfield contributions to memory integration and inference. Hippocampus 24, 1248–1260 (2014).

33. Hitti, F. L. & Siegelbaum, S. A. The hippocampal CA2 region is essential for social memory. Nature 508, 88–92 (2014).

34. Chen, S. et al. A hypothalamic novelty signal modulates hippocampal memory. Nature 586, 270–274 (2020).

35. Lopez-Rojas, J., de Solis, C. A., Leroy, F., Kandel, E. R. & Siegelbaum, S. A. A direct lateral entorhinal cortex to hippocampal CA2 circuit conveys social information required for social memory. Neuron 110, 1559–1572.e4 (2022).

36. Okuyama, T., Kitamura, T., Roy, D. S., Itohara, S. & Tonegawa, S. Ventral CA1 neurons store social memory. Science 353, 1536–1541 (2016).

37. Gilboa, A. & Marlatte, H. Neurobiology of Schemas and Schema-Mediated Memory. Trends Cogn Sci 21, 618–631 (2017).

38. Pliner, P. Cognitive schemas: how can we use them to improve children’s acceptance of diverse and unfamiliar foods? British Journal of Nutrition 99, S2–S6 (2008).

39. Tse, D. et al. Schema-Dependent Gene Activation and Memory Encoding in Neocortex. Science 333, 891–895 (2011).

40. Tse, D. et al. Schemas and Memory Consolidation. Science (1979) 316, 76–82 (2007).

41. Nigro, G. & Neisser, U. Point of view in personal memories. Cogn Psychol 15, 467– 482 (1983).

42. Iriye, H. & St. Jacques, P. L. How visual perspective influences the spatiotemporal dynamics of autobiographical memory retrieval. Cortex 129, 464–475 (2020).

43. Bergouignan, L., Nyberg, L. & Ehrsson, H. H. Out-of-body–induced hippocampal amnesia. Proceedings of the National Academy of Sciences 111, 4421–4426 (2014).

44. Tulving, E. Episodic and semantic memory. in Organization of memory. 423, xiii, 423– xiii (Academic Press, 1972).

45. O’Kane, G., Kensinger, E. A. & Corkin, S. Evidence for semantic learning in profound amnesia: An investigation with patient H.M. Hippocampus 14, 417–425 (2004).

46. Mishkin, M., Vargha-Khadem, F. & Gadian, D. G. Amnesia and the organization of the hippocampal system. Hippocampus 8, 212–216 (1998).

47. Squire, L. R. & Zola, S. M. Episodic memory, semantic memory, and amnesia. Hippocampus 8, 205–211 (1998).

48. Frankland, P. W. & Bontempi, B. The organization of recent and remote memories. Nat Rev Neurosci 6, 119–130 (2005).

49. Tonegawa, S., Morrissey, M. D. & Kitamura, T. The role of engram cells in the systems consolidation of memory. Nat Rev Neurosci 19, 485–498 (2018).

50. Yamanaka, O. & Takeuchi, R. UMATracker: An intuitive image-based tracking platform. Journal of Experimental Biology 221, 1–5 (2018).

