## Supplemental information for "Social transmission of valence-linked new knowledge without firsthand experience in mice"

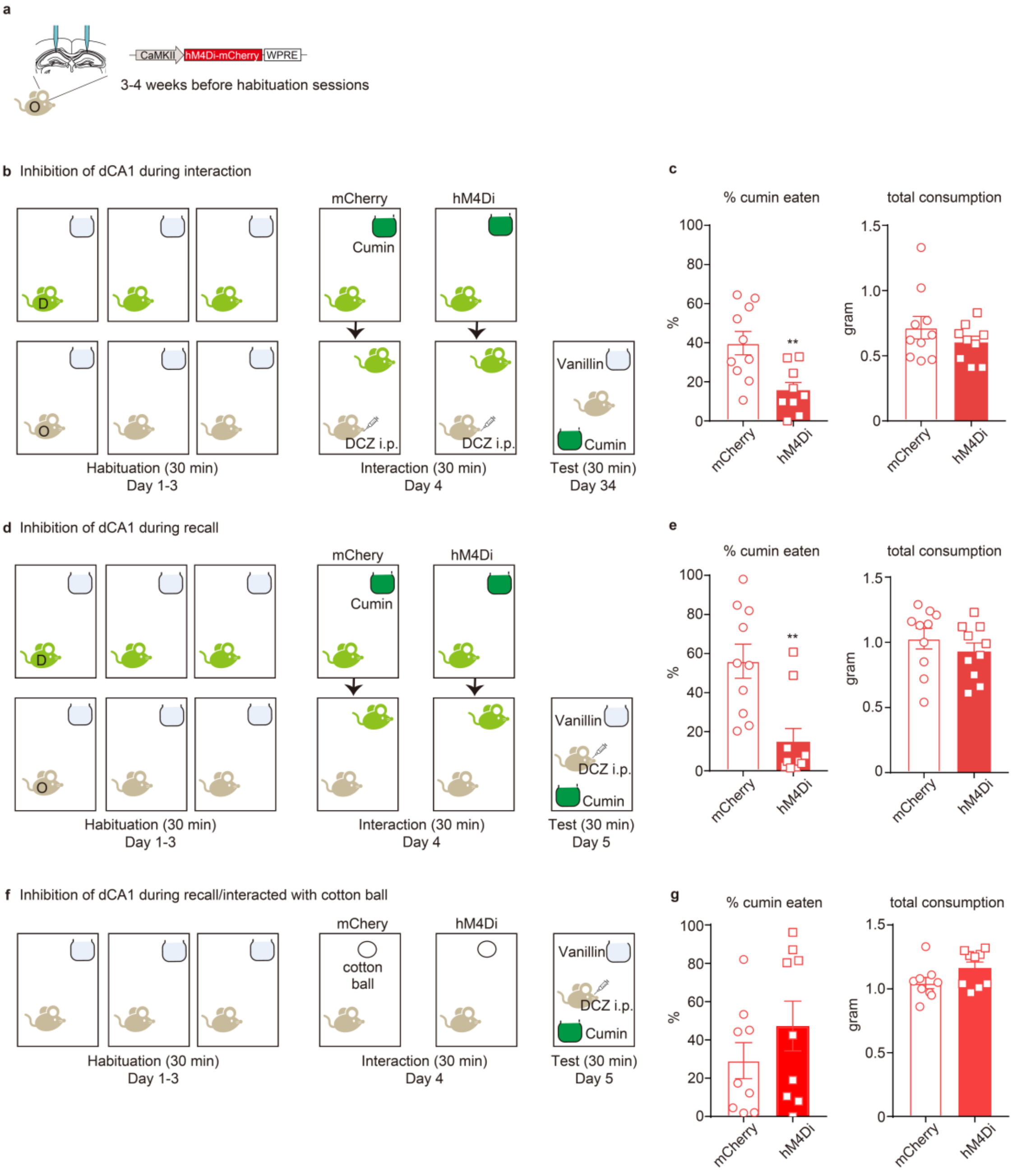


**Supplemental Fig. 1: The dorsal hippocampal CA1 (dCA1) was required for both formation and retrieval of STFP memory in mice, but not in the case of interacting with cotton ball.**

**(a)** The observer mice were injected with hM4Di- or mCherry-expressing AAV into the dCA1. **(b)** Schematic illustration of the protocol for analyzing the role of dCA1 in the formation of STFP memory. D, demonstrator; O, observer. **(c)** The results during the test session. mCherry, n = 10; hM4Di, n = 9. % cumin eaten (Left: two-tailed unpaired *t*-test, $t_{17}$ = 3.278, **P = 0.0044) and total consumption (Right: two-tailed unpaired *t*-test, $t_{17}$ = 1.122, P = 0.2774). **(d)** Schematic illustration of the protocol for analyzing the role of the dCA1 in the retrieval of STFP memory. D, demonstrator; O, observer. **(e)** The results during the test session. n = 10, each group. % cumin eaten (Left: two-tailed unpaired *t*-test, $t_{18}$ = 3.734, **P = 0.0015) and total consumption (Right: two-tailed unpaired *t*-test, $t_{18}$ = 0.9655, P = 0.3471). **(f)** Schematic illustration of the protocol for analyzing the role of the dCA1 in the retrieval of STFP memory. In this case, observer mice interacted with cotton ball instead of demonstrator mice. **(g)** The results during the test session. n = 9, each group. % cumin eaten (Left: two-tailed unpaired *t*-test, $t_{16}$ = 1.124, P = 0.2778) and total consumption (Right: two-tailed unpaired *t*-test, $t_{16}$ = 1.806, P = 0.0898).

**Supplemental table1: Weight of observer mice before the test session.**

| Figure number | | Group | weight (g) | ±SEM | statistical analysis | |
|---|---|---|---|---|---|---|
| Fig. 1 | b-c | Novel | 82.73662017 | 1.081917714 | unpaired *t*-test (two-tailed) | $t_{18}$ = 1.362, p = 0.1899 |
| | | Familiar | 80.82545199 | 0.892854463 | | |
| | e-f | Without | 81.57592219 | 0.774084574 | unpaired *t*-test (two-tailed) | $t_{18}$ = 0.3758, p = 0.7115 |
| | | With | 82.15236072 | 1.324373447 | | |
| Fig. 2 | c | No Cued | 78.42274232 | 0.794471175 | unpaired *t*-test (two-tailed) | $t_{14}$ = 0.4426, p = 0.6648 |
| | | Cued | 78.20396904 | 0.714633682 | | |
| | e | No Cued | 78.42274232 | 0.794471175 | unpaired *t*-test (two-tailed) | $t_{14}$ = 0.4426, p = 0.6648 |
| | | Cued | 78.20396904 | 0.714633682 | | |
| | g | No Cued | 79.17158175 | 0.586323386 | unpaired *t*-test (two-tailed) | $t_{18}$ = 0.4328, p = 0.6703 |
| | | Cued | 78.81565654 | 0.576600768 | | |
| | i | No Cued | 79.26633752 | 0.64434009 | unpaired *t*-test (two-tailed) | $t_{16}$ = 0.9064, p = 0.3782 |
| | | Cued | 78.41996438 | 0.656181432 | | |
| Fig. 3 | b-e | No interaction | 78.93941287 | 0.513245219 | one-way ANOVA | $F_{3,35}$ = 0.8804, p = 0.4606 |
| | | No Odor | 78.8137873 | 0.688212888 | | |
| | | Cumin | 78.11514797 | 0.67018326 | | |
| | | Vanillin | 77.60965351 | 0.770019536 | | |
| Fig. 4 | b | No Odor | 78.91404751 | 0.697355504 | one-way ANOVA | $F_{3,33}$ = 0.7204, p = 0.5469 |
| | | $CS_2$ | 79.98738762 | 0.986141759 | | |
| | | Vanillin | 79.56049116 | 0.508474313 | | |
| | | Vanillin/$CS_2$ | 79.17616564 | 0.704379856 | | |
| | d | No Odor | 78.71161549 | 0.71559564 | unpaired *t*-test (two-tailed) | $t_{18}$ = 1.49, p = 0.1537 |
| | | Vanillin | 79.95265607 | 0.426697645 | | |
| | e | No Odor | 78.26669596 | 0.986142768 | unpaired *t*-test (two-tailed) | $t_{16}$ = 0.4777, p = 0.6393 |
| | | Vanillin | 78.9675139 | 1.086119276 | | |
| Fig. 5 | b | No Odor | 81.62633573 | 1.417861651 | one-way ANOVA | $F_{2,33}$ = 0.05488, p = 0.9467 |

| | | | | | | |
|---|---|---|---|---|---|---|
| | | Saline | 81.03943696 | 1.08270786 | | |
| | | PTZ | 81.50981036 | 1.44753483 | | |
| Fig. 6 | d | VEH-No Cued | 80.32891729 | 1.235678333 | unpaired *t*-test (two-tailed) | $t_{18}$ = 0.07995, p = 0.9372 |
| | | VEH-Cued | 80.2069932 | 0.893691997 | | |
| | | IBO-No Cued | 80.36856347 | 0.875153288 | unpaired *t*-test (two-tailed) | $t_{18}$ = 0.3475, p = 0.7323 |
| | | IBO-Cued | 79.8851885 | 1.081366778 | | |
| | f | Cre only-No Cued | 78.15233532 | 0.930711453 | unpaired *t*-test (two-tailed) | $t_{15}$ = 0.2029, p = 0.8419 |
| | | Cre only-Cued | 77.9136728 | 0.739498264 | | |
| | | tacaspase3-No Cued | 78.03602845 | 0.86513297 | unpaired *t*-test (two-tailed) | $t_{13}$ = 0.532, p = 0.6037 |
| | | tacaspase3-Cued | 78.61740461 | 0.617991319 | | |
| | i | mCherry-No Cued | 77.33915597 | 0.685006046 | unpaired *t*-test (two-tailed) | $t_{16}$ = 0.934, p = 0.3642 |
| | | mCherry-Cued | 78.45096549 | 0.909777144 | | |
| | | hM4Di-No Cued | 78.57826976 | 0.983930167 | unpaired *t*-test (two-tailed) | $t_{16}$ = 0.564, p = 0.5806 |
| | | hM4Di-Cued | 77.99309953 | 0.329221291 | | |
| Supplemental Fig, 1 | c | mCherry | 82.05365806 | 0.897661155 | unpaired *t*-test (two-tailed) | $t_{17}$ = 0.2897, p = 0.7756 |
| | | hM4Di | 81.70109813 | 0.80604092 | | |
| | e | mCherry | 79.83421796 | 0.854623973 | unpaired *t*-test (two-tailed) | $t_{18}$ = 0.3512, p = 0.7295 |
| | | hM4Di | 79.43022695 | 0.769991111 | | |
| | g | mCherry | 80.16380199 | 0.722356941 | unpaired *t*-test (two-tailed) | $t_{16}$ = 1.277, p = 0.2199 |
| | | hM4Di | 78.78840613 | 0.799203651 | | |

**Supplemental video1: Representative behavior during the test session.** Showing mouse represents vanillin group. Chewing behavior was defined as the starting eating the food in the jar. Also see methods.
